# Successful aggregation of Tau protein labelled on its native cysteines

**DOI:** 10.1101/211904

**Authors:** Martina Radli, Romy E Verdonschot, Luca Ferrari, Stefan GD Rüdiger

## Abstract

The formation of fibrillar tangles of the Tau protein is crucial in the development of Alzheimer’s disease. Biophysical methods based on labelling of the cysteines of Tau with fluorescence dyes would allow to study fibril formation with an ‘internal eye’. However, the two native cysteines of Tau at positions 291 and 322 are located in the repeat domain, which is involved in forming the fibrils. The contribution of both cysteines to this process is unclear. Here we show that blocking natural cysteines using large fluorescent dyes does not interfere with Tau fibrillation so that FRET can be used to follow structural changes during the process. We anticipate that cysteine-labelled Tau enables following structural rearrangements during fibril formation in detail. This may also allow to monitor the effect of drugs, small molecules and proteins on the process.

## Introduction

The intrinsically disordered Tau protein is involved in stabilisation and organisation of axonal microtubule bundles due to its microtubule-binding and tubulin-polymerising activity ^1, 2^. Tau is abundant in neurons, especially in axons and its aggregation into paired helical filaments and neurofibrillary tangles are the key hallmarks of multiple neurodegenerative disorders, including Alzheimer’s disease (AD) and other tauopathies ^3–6^. Notably, the literature refers to fibril formation of Tau as aggregation. However, when Tau aggregates it always assembles into structurally defined fibrils, Tau does not form amorphous aggregates.

Aggregation enhancers facilitate fibril formation of Tau and a variety of methods allow monitoring fibrils. As the first step of the process the structural ensemble of Tau undergoes significant transitions which leads to oligomers of Tau ^7^. Subsequently, oligomers are elongated and form long, insoluble fibrils ^8^. During fibril formation the highly soluble, unfolded Tau enriches in β-sheet elements and adopts a combined cross-β/β-helix structure ^9–11^. Recent studies suggest that oligomeric Tau species, rather than fibrils, are responsible for cellular toxicity and disease pathology. However, the mechanism of oligomer toxicity is poorly understood ^12–20^.

Tau consists of two major domains of which the repeat domain (Tau-RD) in the assembly domain primarily drives fibril formation (**Figure 1A**) ^21^. It is not yet known how the projection domain and the connecting proline rich region influence polymerisation. However, the termini of Tau are suggested to have an aggregation inhibitory effect by folding on the Tau-RD and preventing its self-association ^22, 23^. The Tau-RD, depending on alternative splicing, consists of three or four pseudorepeats that form the core of the paired helical filaments upon Tau aggregation ^21, 24, 25^. The minimal motifs of the Tau-RD that were shown to be necessary for Tau aggregation *in vitro* are the (^275^VQIINK^280^) and (^306^VQIVYK^311^) hexapeptides, located in the second and third repeats, respectively (**Figure 1B**) ^26^. A recent high resolution Cryo-EM structural model of Tau paired helical filaments from an Alzheimer’s patient suggests that the filament core consists of two identical copies comprising the residues of V306-F378 ^9^. Additionally, a section N-terminally from the core folds back forming a less ordered β-sheet with the filament core.

**Figure 1:**
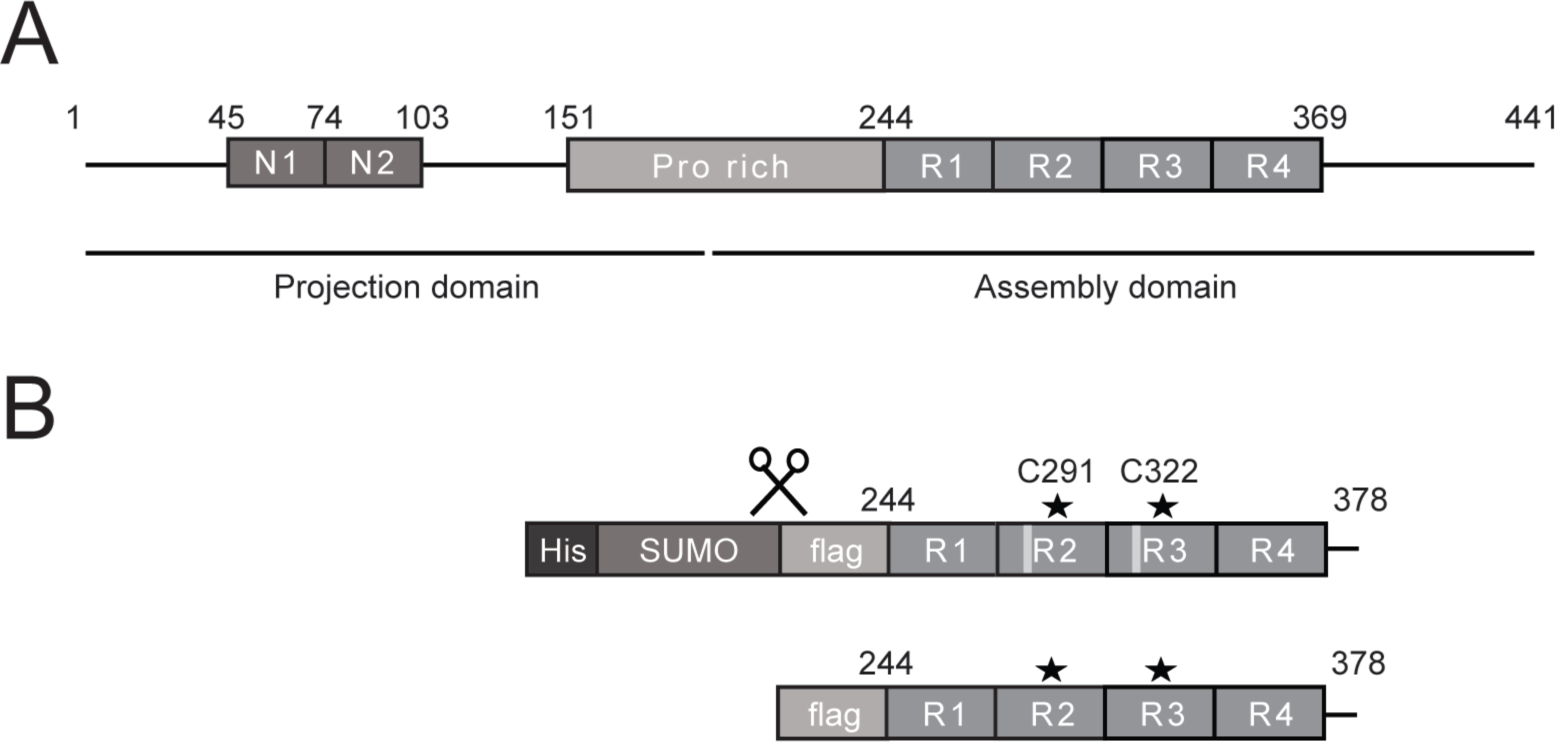
Tau(Q244-E372) is sufficient for aggregation. (A) Domain structure of Full length Tau protein. The projection domain consists of 0, 1 or 2 near N-terminal inserts (N1 and N2) and part of the proline rich connecting region. The other part of the proline rich region and 3 or 4 pseudo-repeats (R1-R4, R2 may be spliced out) make up the C-terminal microtubule assembly domain. The pseudo-repeats are sufficient for microtubule assembly and aggregation. (B) The Tau(Q244-E372) construct used in this study consists of the four pseudo-repeats, an N-terminal flag-tag and a short C-terminal flanking. Initially it is produced with a His_6_-SUMO-tag that is cleaved during the purification process.

Tau has two native cysteine residues, one located in the filament core and the other on the region that folds back on the core (**Figure 1B**) ^9^. It is controversial to which extent they contribute to or even drive aggregation. It was shown that Tau molecules form covalent dimers via intermolecular disulfide bridges between their free native cysteines in oxidative environment. This is suggested to be essential for aggregation initiation since constructs without cysteines failed to drive the process ^27, 28^. Besides, intramolecular disulfide bond formation by cysteine over-oxidation is aggregation-inhibitory, presumably because of the lack of free cysteines ^29–32^. Furthermore, 1,2-dihydroxybenzene-containing compounds that bind to and ‘cap’ cysteine residues also inhibit aggregation ^33^. Taken together these data suggest that free/accessible cysteines may be important to drive the first steps of aggregation.

In contrast, others were able to aggregate cysteine-free constructs, although morphology and aggregation propagating capacity of the resulting fibrils were different from wild type fibrils ^34–36^. Furthermore, Tau labelled with fluorescent dyes on artificially introduced cysteines in the Tau-RD aggregated, suggesting that the presence of large modifications on cysteines does not inhibit the process ^23^. These data indicate that even though free cysteine side chains may have a role in Tau aggregation, they are not essential.

We set out to understand if labelling the native cysteine residues of Tau has an effect on its aggregation process. Therefore, we labelled aggregation-prone Tau-RD with two dyes forming a FRET-pair and performed aggregation experiments with this sample. We visualised the polymerisation process using blue native-PAGE and in parallel, monitored structural rearrangements by FRET.

We found that the presence of large fluorescent modifications on native cysteines allowed Tau aggregation. The protein formed oligomers and fibrils upon addition of aggregation enhancer. FRET-labels enabled us to follow conformational changes that accompanied the aggregation process. We anticipate that this experimental setup will provide a useful tool to monitor structural changes of aggregating Tau with the possibility to test the effect of additional compounds (proteins, small molecules, drug targets) on the process.

## Results

### Labelling of Tau-RD by fluorescent dyes

In our experiments we used a construct that consists of the four pseudo-repeats (R1-R4) of Tau (Tau-RD), Tau(Q244-E372), also referred to as K18 in the literature. This protein is established to aggregate faster than the full length as the termini of Tau have aggregation-inhibitory effect ^22, 25, 37–39^. We modified Tau(Q244-E372) with an N-terminal flag-tag for western blot detection and a K280 deletion that has pro-aggregating effect ^40^. We labelled the Tau(Q244-E372) fragment using the FRET-pair Alexa 488 (donor) and Alexa 594 (acceptor) dyes with maleimide reaction groups to block cysteine residues (**Figure 1B**).

The absorption spectrum of the double-labelled and purified protein showed two maxima at 495 nm and 594 nm, thus, we concluded that simultaneous labelling of Tau(Q244-E372) with Alexa 488 and Alexa 594 was successful and it resulted in double-labelled Tau(Q244-E372) (Tau(Q244-E372)^*488D*594A^). This suggests that the cysteines of the Tau(Q244-E372) fragment are accessible. In order to calculate the labelling efficiency we measured the label concentration by absorbance spectroscopy and Tau(Q244-E372) concentration using bicinchoninic colorimetry assay (BCA). Since the Tau(Q244-E372) fragment has two cysteines, at 100% labelling efficiency one mole of Tau(Q244-E372) contains two moles of dye. In our experiments, one mole of Tau(Q244-E372) typically contained between 1.4 and 1.8 moles of dye.

We set out to measure whether we could detect FRET signal in the Tau(Q244-E372)^*488D*594A^ sample. We excited Tau(Q244-E372)^*488D*594A^ at 470 nm since at this wavelength only the donor dye has significant absorbance **(Figure 2A**). We monitored fluorescence emission between 590 and 650 nm as at this wavelength interval the donor dye (Alexa 488) has low, while the acceptor dye (Alexa 594) has high emission **(Figure 2A**). We were able to detect FRET signal in the Tau(Q244-E372)^*488D*594A^ sample (**Figure 2B,** black bar).

**Figure 2:**
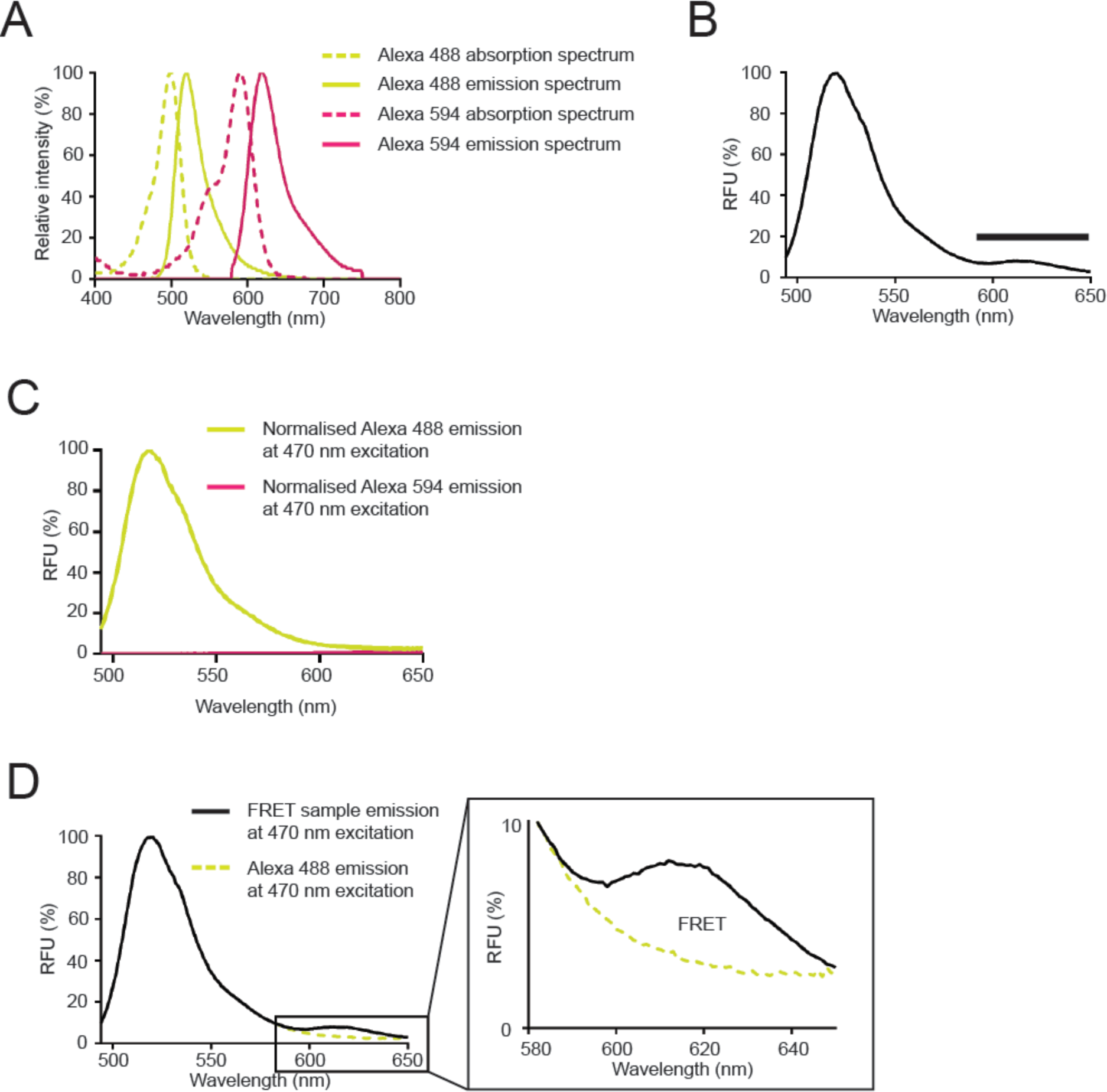
Successful labelling of Tau(Q244-E372) by the FRET-pair Alexa 488 and Alexa 594. (A) Excitation (dotted line) and emission (solid line) spectra of Alexa 488 (donor, yellow line) and Alexa 594 (acceptor, magenta line) normalised to 100%. (B) Emission spectrum of the Tau(Q244-E372)^*488D*594A^ construct at the excitation wavelength of 470 nm. (C) Normalised Alexa 488 (donor, yellow line) and Alexa 594 (acceptor, magenta line) emission spectra at the excitation wavelength of 470 nm. (D) Left panel: Normalised emission spectra of Tau(Q244-E372)^*488D*594A^ (black solid line) and Tau(Q244-E372)^*488D^ (yellow, solid line). Inset: Visualisation of the appearance of the FRET signal in our experimental setup. The difference of the Tau(Q244-E372)^*488D*594A^ and the Tau(Q244-E372)^*488D^ corresponds to the FRET signal.

Since both the donor and the acceptor dyes may have contributed to the measured FRET signal we checked whether we needed to correct for their effect (**Figure 2C)**. Therefore, we labelled Tau(Q244-E372) with Alexa 488 (Tau(Q244-E372)^*488D^) or Alexa 594 (Tau(Q244-E372)^*594A^) and measured the emission spectrum of the samples at 470 nm excitation. The donor dye had considerable emission between 590 and 650 nm upon excitation at 470 nm (**Figure 2C,** yellow). To correct for this effect, we normalised the emission spectra of Tau(Q244-E372)^*488D*594A^ and Tau(Q244-E372)^*488D^ to 100% based on the emission maxima of the spectra and calculated their difference (**Figure 2D** and **Figure 2D** inset). The ratiometric FRET efficiency was ~10% of the total fluorescence signal. Since the acceptor dye has very low absorption at 470 nm and therefore very low emission between 590 and 650 nm, we did not correct for its effect (**Figure 2 C,** magenta).

We concluded that the cysteines of Tau(Q244-E372) are accessible and the protein could be labelled simultaneously using multiple different chemicals with maleimide reactive groups. We detected FRET signal in the Tau(Q244-E372)^*488D*594A^ construct after donor correction, suggesting that the two cysteine residues are within FRET distance from each other.

### intramolecular FRET in cysteine-labelled Tau(Q244-E372)

After successful labelling we set out to understand whether the FRET signal we detected in the Tau(Q244-E372)^*488D*594A^ sample resulted from inter- or intramolecular energy transfer. We excited Tau(Q244-E372)^*488D*594A^ and a mix of Tau(Q244-E372)^*488D^ and Tau(Q244-E372)^*594A^ at 470 nm and compared the fluorescence emission spectra of the samples between 590 nm and 650 nm. We expected to detect FRET in both samples if it originates from intermolecular but only in the Tau(Q244-E372)^*488D*594A^ sample if it is the result of intramolecular FRET.

After normalisation and donor correction of the fluorescence emission of Tau(Q244-E372)^*488D*594A^ we observed a peak between 590 and 650 nm indicating that energy transfer occurred (**Figure 3A**). In contrast, we did not observe any peak in the normalised and donor-corrected emission spectrum of the mix of Tau(Q244-E372)^*488D^ and Tau(Q244-E372)^*594A^ sample between 590 and 650 nm (**Figure 3B**). This suggests that intermolecular interactions did not contribute to the detected FRET signal. We concluded, the FRET signal is a result of intramolecular interactions.

**Figure 3:**
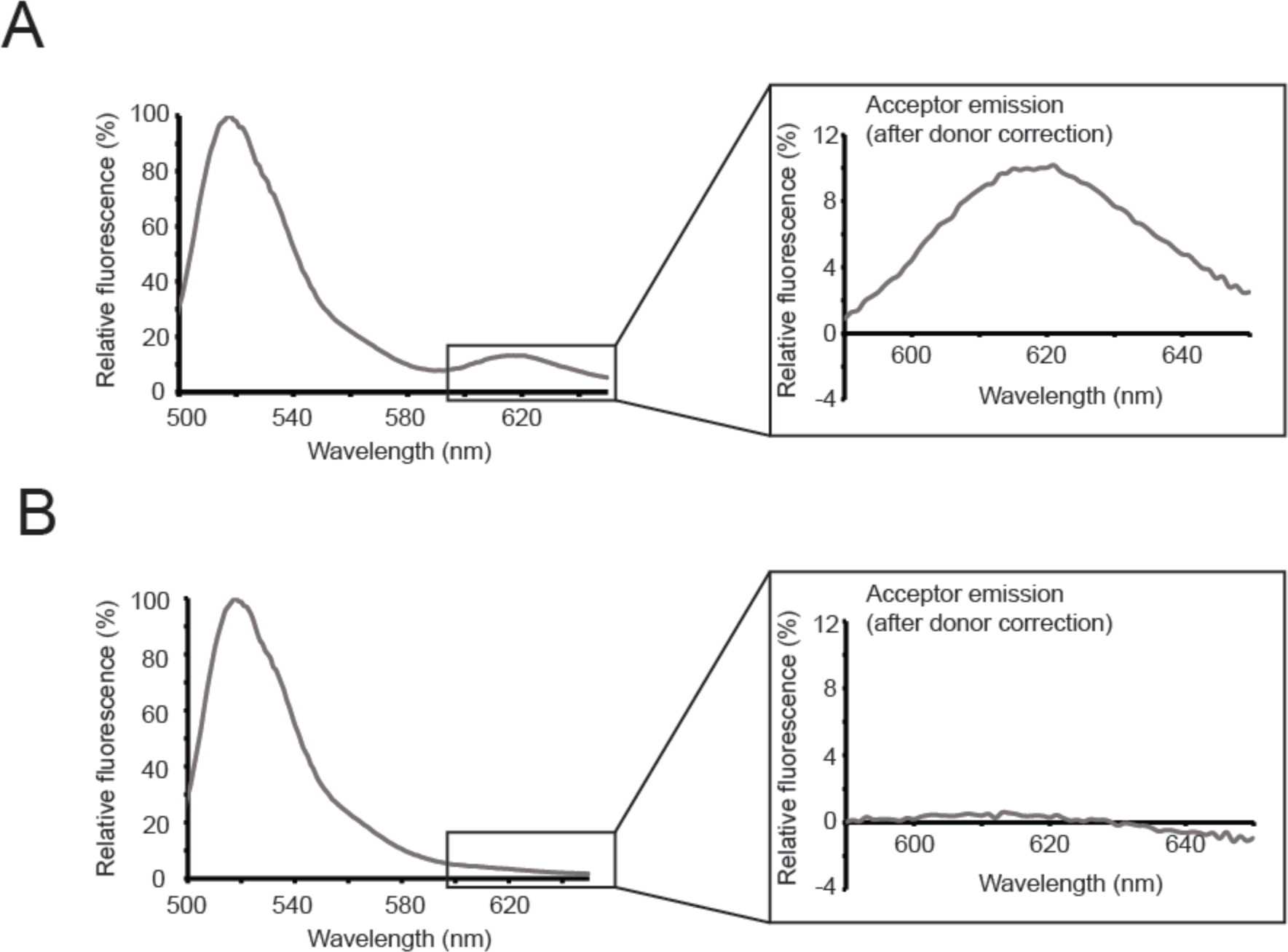
Only intramolecular FRET before aggregation. (A) Left panel: Emission spectrum of Tau(Q244-E372)^*488D*594A^ at the excitation wavelength of 470 nm. Inset: FRET signal of the same sample after donor correction. (B) Left panel: Emission spectrum of the mix of Tau(Q244-E372)^*488D^ and Tau(Q244-E372)^*594A^ at the excitation wavelength of 470 nm. Inset: No FRET signal is detected after donor correction.

### Tau(Q244-E372) labelled on its native cysteines aggregates

We used the Tau(Q244-E372)^*488D*594A^ construct to test whether the reduction in the number of free cysteines and the presence of large cysteine modifications inhibit the aggregation process as suggested earlier ^27, 28^.

We simultaneously aggregated unlabelled Tau(Q244-E372) and Tau(Q244-E372)^*488D*594A^ using heparin, a well-established polyanionic aggregation enhancer of Tau ^28, 30, 41^. We took samples at given time points throughout the experiment to be able to visualise the different molecular weight species that have been forming during Tau(Q244-E372) polymerisation and to gain insight into the kinetics of the process. We used blue native-PAGE to separate those species since non-covalent interactions that stabilise oligomers and fibrils remain intact with this method. After running the samples on blue native gels, we transferred them to a membrane and visualised the monomers, oligomers and fibrils/high molecular weight species by blotting for the flag-tag of Tau(Q244-E372).

In the absence of heparin we detected only monomers and oligomers in the Tau(Q244-E372)^*488D*594A^ sample (**Figure 4A**). The amount of these species did not change over time and we did not observe fibril formation. This is consistent with the fact that Tau(Q244-E372) aggregates slowly without heparin. To the contrary, upon addition of heparin the amount of monomers decreased and eventually disappeared over time in the Tau(Q244-E372)^*488D*594A^ sample (**Figure 4A**). We observed first increase and then decline in oligomer amount and gradual fibril formation. We concluded that we were able to initiate the aggregation of Tau(Q244-E372)^*488D*594A^ with the use of heparin. Consequently, large modifications on the native cysteine residues of Tau(Q244-E372) did not inhibit the aggregation of the protein.

**Figure 4:**
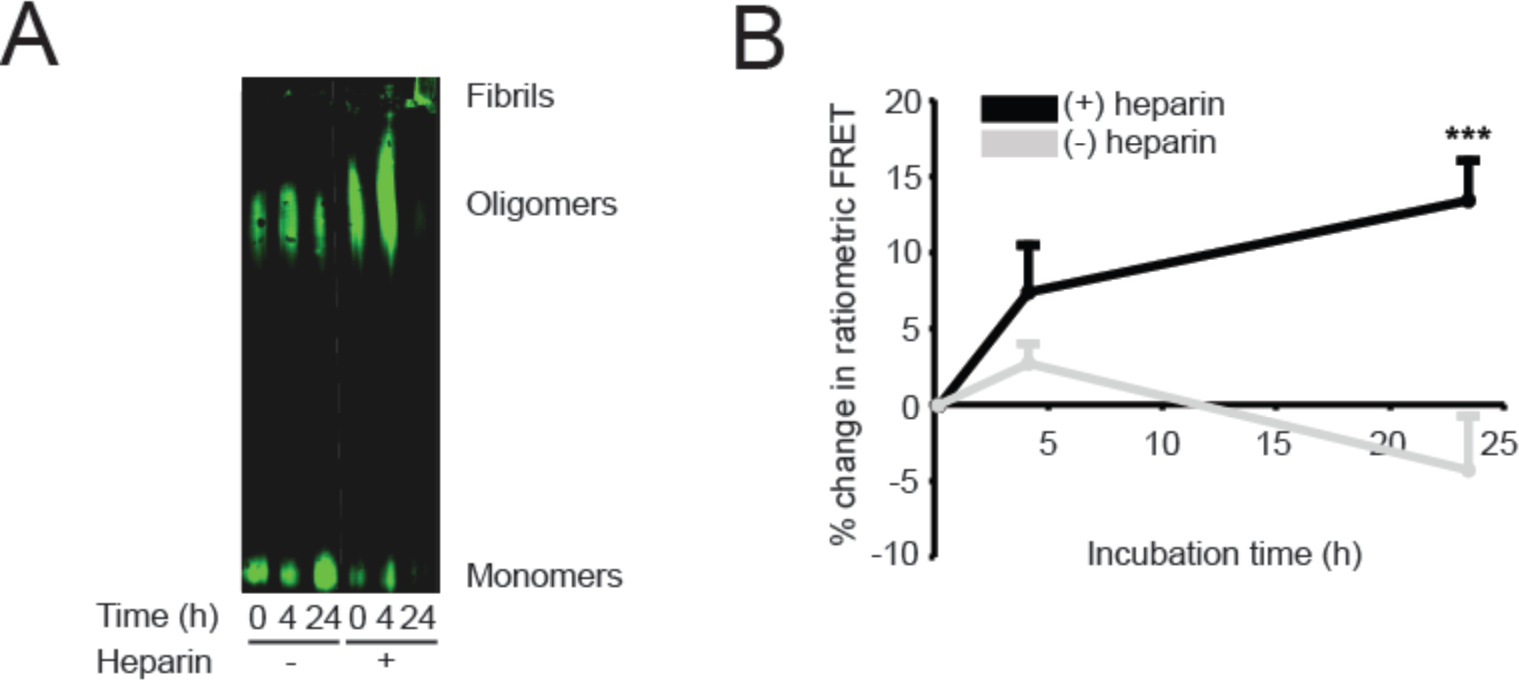
Flag-Tau(Q244-E372) labelled on its native cysteine residues aggregated upon addition of heparin. (A) Western blot of cysteine-labelled Tau(Q244-E372)^*488D*594A^ incubated on 37°C for 0, 4 and 24 hours in the absence (-) or presence (+) of heparin. (B) The ratiometric FRET efficiencies of aggregated Tau(Q244-E372)^*488D*594A^ -samples ((grey: (-) heparin, (n=2), black: (+) heparin (n=6)) were determined after 0, 4 and 24 hours of incubation. Percentage change in FRET (mean ± SEM) is plotted as a function of time, ** < 0.01, *** < 0.001 compared to 0 hours as determined by a two-way ANOVA with LSD-corrected post hoc test.

Next, we set out to test whether the FRET setup allowed us to follow structural rearrangements that accompany aggregation. In the time points of the native gel samples we measured fluorescence emission of Tau(Q244-E372)^*488D*594A^ between 590 and 650 nm at the excitation wavelength of 470 nm. By calculating the percentage change in FRET efficiency we could follow structural rearrangements that accompany aggregation (**Figure 4B**). In the absence of heparin when no aggregation happened ratiometric FRET efficiency did not change over time. In contrast, ratiometric FRET efficiency significantly increased when we aggregated the protein in the presence of heparin, suggesting that compacting accompanied the process.

In summary, we were able to label the repeat domain of Tau using fluorescent dyes that form a FRET-pair. We showed that this construct aggregated in the presence of aggregation enhancer and that FRET was suitable to follow structural rearrangements during the course of aggregation.

## Discussion

We successfully labelled the repeat domain of Tau on its native cysteines using large fluorescent dyes that form a FRET-pair. We detected FRET signal in the double-labelled Tau(Q244-E372)^*488D*594A^ but not in the mix of the single-labelled Tau(Q244-E372)^*488D^ and Tau(Q244-E372)^*594A^ samples. This suggests that the energy transfer between both dyes of the FRET pair was intramolecular.

According to our results, labelling Tau on its native cysteines enables the aggregation of the protein. This is in contrast with some earlier findings, since the necessity of free cysteines for aggregation was strongly suggested previously ^27–32^. As the labelling efficiencies of our samples were between 70 and 90%, they contained a subpopulation with free cysteines. Therefore, in principle dimer formation via intermolecular disulfide bonds as the primary step of the aggregation could have occurred in our samples. However, we detected oligomerisation and fibrillation in the presence of high concentration of reducing agents, which substantially reduced the chance of covalent dimer formation. Furthermore, we never observed dimers on native gels (**Figure 4A**). Successful aggregation experiments with cysteine-free constructs also argue that covalent dimers are not essential for the process, however, aggregation dynamics and fibril morphology was affected by the absence of cysteines ^34, 35^. The differences between these results may originate from variations in the experimental conditions (especially the presence or absence of reducing agents) and the type of modifications.

The presence of large modifications on the native cysteines of Tau did not prevent aggregation in our experiments. This was surprising, because one would expect that bulky dyes inhibited the tight packing of Tau fibrils. Furthermore, it has been shown that shielding natural cysteines by 1,2-dihydroxybenzene-containing compounds has aggregation inhibitory effect ^33^. A recent high resolution Cryo-EM structure of full length and truncated Tau aggregates derived from an AD patient, however, significantly improved our understanding of the three dimensional conformation of Tau paired helical filaments ^9^. According to this structure Tau adopts a combined cross-β/β-helix conformation where C322 is part of the filament core, whereas C291 is in the less ordered β-sheet that folds on the core. Both cysteine residues seem to be accessible, as the authors did not find regular density hinting towards intermolecular disulfide bonds between C322 residues and C291 is located in a relatively disordered region ^9^. This suggests that despite the compactness of Tau tangles there is enough room to accommodate modifications covalently attached to Tau’s native cysteines. Furthermore, similarly to our experiments labelling of non-natural cysteines by fluorescent dyes enabled aggregation ^23^. We concluded that the presence of fluorescent dyes seem to allow the progression of aggregation regardless of their positions.

## Experimental procedures

### The ΔK280 (pro-aggregating) His_6_-SUMO-flag-Tau(Q244-E372) construct

pET SUMO expression vector, a kind present from Elke Dauerling, was used for the recombinant production of human His_6_-SUMO-flag-tagged ΔK280 (pro-aggregating) Tau(Q244-E372) construct in Rosetta 2 *E. coli* strain (Novagen). The DNA construct was cloned by Luca Ferrari. The sequence of the construct: MGHHHHHHGSDSEVNQEAKPEVKPEVKPETHINLKVSDGSSEIFFKIKKTTPLRRLMEAFAKRQGKEMDSLRFLYDGIRIQADQTPEDLDMEDNDIIEAHREQIGGDYKDDDDKMQTAPVPMPDLKNVKSKIGSTENLKHQPGGGKVQIINKLDLSNVQSKCGSKDNIKHVPGGGSVQIVYKPVDLSKVTSKCGSLGNIHHKPGGGQVEVKSEKLDFKDRVQSKIGSLDNITHVPGGGNKKIE

### Production of ΔK280 His_6_-SUMO-flag-Tau(Q244-E372)

Rosetta 2 cells containing pET SUMO vector with the His_6_-SUMO-flag-ΔK280-Tau(Q244-E372) (Figure 1) were inoculated into 100 ml 2xYT (12.8 g Bacto Tryptone (Merck), 8 g Bacto Yeast Extract (Merck), 4 g NaCl (Merck) in 800 ml demi water (Millipore) medium supplemented with 34 mg/l final concentration of chloramphenicol (Sigma-Aldrich) and 50 mg/l final concentration of kanamycin (Sigma-Aldrich) (final concentration). The cells were grown over night on 37°C, shaking with 180 rpm. Next morning the 100 ml culture was inoculated into 4×800 ml 2xYT medium supplemented with 34 mg/l final concentration of chloramphenicol and 50 mg/l final concentration of kanamycin. The cultures were grown on 37°C, shaking with 180 rpm.

The OD_600_ was monitored every 1.5 hours by measuring the mix of 900 µl medium and 100 µl cell culture by Ultrospec 3000 pro UV/Visible Spectrophotometer (GE Healthcare).

The cultures were induced at the OD_600_ of 0.8-1.0 with 0.2 mM IPTG (Thermo Scientific), and incubated at 18°C overnight. The cultures were harvested in an Avanti J-26 XP centrifuge (Beckman Coulter) using the JLA-8.1 rotor on 4°C at 4500 rpm for 30 minutes. The supernatant was discarded and the pellet was resuspended in ice cold resuspension-buffer (50 mM Na-phosphate pH 7.2 (Sigma-Aldrich), 150 mM NaCl, 150 mM KCl (CARL ROTH)) and centrifuged in an MSE Harrier 18/80 Refrigerated Benchtop Centrifuge on 4°C at 5000 rpm for 30 minutes. The supernatant was discarded and the pellet was stored on −20°C until further usage.

### Purification of ΔK280 His_6_-SUMO-flag-Tau(Q244-E372)

The pellet was resuspended in ice cold lysis buffer (25 mM Tris pH 8.5 (Sigma-Aldrich), 20 mM NaCl, 5 mM β-mercaptoethanol (Sigma-Aldrich), EDTA-free protease inhibitor (1 tablet/50 ml) (Roche)). The cells were disrupted by an EmulsiFlex-C5 (Avestin) cell disruptor. The lysate was centrifuged in Avanti J-26 XP centrifuge using JA-25.5 rotor at 21,000 rpm for 45 minutes on 4°C. The lysate was filtered by 25 mm 22 µm polypropylene filter (VWR) to remove the cell debris and insoluble aggregates. The purification was done using an AKTA Purifier (GE Healthcare).

His_6_-SUMO-flag-ΔK280-Tau(Q244-E372) was first purified on an IMAC POROS 20MC (Thermo Fischer Scientific) affinity purification column (solutions connected to pump A1 and A2: 50 mM Na-phosphate buffer pH 8.0, B1: demi water with 10 mM β-mercaptoethanol, B2: 1 M imidazole (Sigma-Aldrich)). The elution was concentrated using a Vivaspin 20 column (5 kDa MWCO) (GE Healthcare) on 4°C at 5000 rpm for 15-15 minutes in a MSE Harrier 18/80 Refrigerated Benchtop Centrifuge until we reached 2.5 ml volume. The concentrated sample was incubated overnight on 4°C with Ulp1 SUMO-protease to cleave the His_6_-SUMO-tag.

Next day the sample was buffer exchanged with a PD10 desalting column (GE Healthcare) to 50 mM Tris pH 8.0 buffer supplemented with 5 mM β-mercaptoethanol and protease inhibitor (1 tablet/50 ml) (Roche, Complete). To separate flag-Tau(Q244-E372) from the His_6_-SUMO-tag, the protein sample was loaded on a POROS 20HS cation exchange column (Thermo Fischer Scientific) (solutions connected to pump A1 and A2: 50 mM Na-phosphate pH 7.2, B1: demi water with 10 mM DTT (Sigma-Aldrich), B2: 2 M KCl). The elution was frozen in liquid N_2_ and stored at −80°C until further usage. Protein purity was determined with SDS-PAGE, protein concentration was measured with a Pierce BCA Protein Assay Kit (Thermo Fisher Scientific) according to the manufacturer’s instructions.

### Fluorescent labelling of the ΔK280 flag-Tau(Q244-E372)

Purified flag-ΔK280-Tau(Q244-E372) was 100x buffer-exchanged to labelling buffer (50 mM HEPES pH 7.2 (Sigma-Aldrich), 150 mM NaCl 1 mM TCEP (Sigma-Aldrich), protease inhibitor (1 tablet/50 ml)) using a Vivaspin 6 column (5 kDa MWCO) on 4°C at 5000 rpm for 15-15 minutes in a MSE Harrier 18/80 Refrigerated Benchtop Centrifuge. The protein was incubated overnight at 4°C with a 10-fold molar excess mix of Alexa 488 and 594-maleimide fluorescent dyes (Thermo Fisher Scientific). Additionally, flag- ΔK280-Tau(Q244-E372) was labelled with a mix of N-Ethylmaleimide (non-fluorescent compound with maleimide reactive group) (Sigma-Aldrich) and either Alexa 488 or Alexa 594 for single-cysteine fluorescent labelling using a similar procedure. Labelled flag-ΔK280-Tau(Q244-E372) was buffer-exchanged to aggregation buffer (25 mM HEPES pH 7.5, 75 mM NaCl, 150 mM KCl, 10 mM DTT and protease inhibitor (1 tablet/50 ml)) using a PD MiniTrap G-25 desalting column, according to the instructions of the manufacturer. Subsequently, labelled flag-ΔK280-Tau(Q244-E372) was concentrated and isolated from free dye using an Amicon Ultra-0.5 ml Centrifugal Filter (5 kDa MWCO) (Millipore) by repeated centrifugation cycles of 5 min at 4500 rpm. The concentration of the protein was determined by SDS PAGE using a bovine serum albumin (BSA) standard and with a Pierce BCA Protein Assay Kit (Thermo Fisher Scientific) according to the manufacturer’s instructions. The concentration of the labels was determined with ND-1000 program on an ND-1000 Spectrophotometer type NanoDrop after which labelling efficiency was determined.

### Concentration determination by SDS-PAGE

For SDS-PAGE BSA (Sigma-Aldrich) standard was prepared, 1:1 mixed with 2x sample buffer (0.625 M Tris, 12.5% glycerol (CARL ROTH), 1% SDS (Bio-Rad), 0.005% Bromophenol Blue (Bio-Rad), 5 mM freshly added β-mercaptoethanol) and was loaded on the gel in a strictly determined amount. Labelled flag-ΔK280-Tau(Q244-E372) samples were 1:1 mixed with 2x sample buffer (buffer (0.625 M Tris, 12.5% glycerol, 1% SDS, 0.005% Bromophenol Blue, 5 mM freshly added β-mercaptoethanol) and together with the BSA standard, they were loaded on the SDS-PAGE.

15% SDS gels were prepared (Separation buffer: 0.38 M Tris pH 8.8, 15% acrylamide (National Diagnostics), 0.1% SDS, 0.1% APS (Sigma-Aldrich), 0.04% TEMED (Sigma-Aldrich), Stacking buffer: 0.125 M Tris pH 6.8, 4% acrylamide, 0.1% SDS, 0.075% APS, 0.1% TEMED) and ran in 1x Laemmli buffer (0.025 M Tris base, 0.152 M glycine (SERVA Electrophoresis GmbH), 0.1% SDS, diluted from 10x stock). The gels were stained with Coomassie staining solution (0.2% Coomassie Brilliant Blue (SERVA Electrophoresis GmbH), 45% methanol (Interchema Antonides-Interchema), 10% acetic acid (Biosolve) and 55% demi water) and destained by destaining solution (30% methanol, 10% acetic acid and 60% demi water).

The density of the flag-ΔK280-Tau(Q244-E372) band was compared to the density of the BSA bands and the concentration of flag-ΔK280-Tau(Q244-E372) was estimated. This was needed because there is a concern that the fluorescent labels interfere with the signal of the BCA Assay.

### Aggregation of flag-Tau(Q244-E372)

For flag-Tau(Q244-E372) aggregation, a 150 µl aggregation mix was prepared in aggregation buffer (25 mM HEPES pH 7.5, 75 mM NaCl, 150 mM KCl, 10 mM DTT and protease inhibitor (1 tablet/50 ml)). The aggregation mix consisted of 10 µM flag-Tau(Q244-E372), protease inhibitor, 10 mM DTT, 4/1 molar ratio (protein/heparin) heparin (Santa Cruz Biotechnology) and 10-10 µM of protein (Hsp90) when added. 100 µl of the mix was added to a well of a 96-well plate (black, Corning) for FRET measurements and 50 µl of the mix was added to separate well for native gel samples. The plate was covered, sealed with parafilm and incubated on 37°C shaking with 180 rpm for 24 hours. At 0, 4 and 24 hours the FRET signal of the samples was measured by exciting the sample with a wavelength of 470 nm and recording the emission from 580 till 650 nm. Simultaneously 10 µl gel samples were taken, snap-frozen in liquid nitrogen and stored at −20°C overnight.

### FRET

Fluorescence measurements were performed with a CLARIOstar^®^ microplate reader (BMG Labtech). Samples containing 10 µM labelled Tau(Q244-E372) with the volumes of 100 µl were excited at 470 nm to prevent direct excitation of the acceptor (Alexa 594). Fluorescence was measured from 494 to 650 nm. To control for the spectral overlap of the FRET pair, donor (D) (Tau(Q244-E372)^*488^) fluorescence in absence of acceptor (A) and acceptor (Tau(Q244-E372)^*594^) fluorescence in absence of donor were measured and the donor fluorescence was normalised. Next, the fluorescence signal of double-labelled sample (Tau(Q244-E372)^*488*594^) was also measured. The fluorescence signal of Tau(Q244-E372)^*488*594^ was normalised and corrected by D fluorescence by deducting D fluorescence from Tau(Q244-E372)^*488*594^ fluorescence. The ratiometric FRET efficiency was then calculated by the equation: I_A_ / (I_D_ + I_A_), where I_A_ is the intensity of donor-corrected Tau(Q244-E372)^*488*594^ fluorescence at the emission wavelength of the acceptor, I_D_ is the intensity of Tau(Q244-E372)^*488*594^ fluorescence at the emission wavelength of the donor ^42^.

### Western blot

The aggregation samples were thawed on ice and 1 µl of sample buffer (0.1% Ponceau S (Sigma-Aldrich), 50% glycerol) was added to them. The samples were ran on blue native-PAGE (BN-PAGE) ^43^ using 4~16% Bis-Tris gradient gels (Novex), following the instructions of the manufacturer. Fibrils and large aggregated species above 1500 kDa remained in the stacking part, whereas oligomers and monomers migrated into the separation part of the gel. Proteins were transferred to PVDF membranes (0.45 µm pore size) (Immobilon) overnight at 4°C in transfer buffer (25 mM Tris, 192 mM glycine and 20% methanol) at constant current (10 mA/gel). Next day membranes first were washed with methanol three times for 2 min, rinsed with demi water, washed with TBS (50 mM Tris pH 7.6, 150 mM NaCl) for 2 min and blocked with TBS Odyssey blocking buffer (LI-COR) for 1 hour at room temperature. The membranes were incubated in TBS-T ((50 mM Tris pH 7.6, 150 mM NaCl, 0.1% Tween-20 (Sigma-Aldrich)) with a 1:500 dilution of mouse α-FLAG monoclonal primary antibody (F1804, Sigma) for 1 hour at room temperature, then the blots were washed four times for 5 min with TBS-T. Subsequently the membranes were incubated in TBS-T with 1:1000 dilution of IRDye 800CW Donkey anti-Mouse IgG antibody (LICOR) or IRDye 680RD Goat anti-Rabbit IgG antibody (LICOR) secondary antibody for 1 hour at room temperature, then the blots were washed four times for 5 min with TBS-T. The membrane was washed two times with TBS-T and two times with TBS prior to detection using the Odyssey scanner (LI-COR).

## Acknowledgements

We are grateful to Ineke Braakman for continuous support. SGDR was supported by Marie-Curie Actions of the 7th Framework programme of the EU [Innovative Doctoral Programme “ManiFold” (No. 317371) and Initial Training Network “WntsApp” (No. 608180)], the Internationale Stichting Alzheimer Onderzoek (ISAO; project “Chaperoning Tau Aggregation”; No. 14542) and a ZonMW TOP grant (“Chaperoning Axonal Transport in neurodegenerative disease”; No. 91215084).

## Author contributions

The concept of the study originates from M. Radli and S.G.D. Rüdiger. M. Radli drafted the first version of the manuscript and prepared figures 1 and 3. R.E. Verdonschot provided the images for figures 2 and 4. M. Radli and S.G.D. Rüdiger wrote the manuscript.

M. Radli produced, purified and labelled Tau(Q244-E372) and performed the experiments to check for inter- and intramolecular FRET under the supervision of S.G.D. Rüdiger. R.E. Verdonschot produced, purified and labelled Tau(Q244-E372). She calculated the values for donor correction, performed aggregation experiments with double-labelled Tau(Q244-E372), ran the BN-PAGE gels and calculated the FRET efficiencies under the supervision of M Radli and S.G.D. Rüdiger. L. Ferrari cloned the Tau(Q244-E372) fragment and established the aggregation experiment under the supervision of S.G.D. Rüdiger.

